# A metric and its derived protein network for evaluation of ortholog database inconsistency

**DOI:** 10.1101/2022.01.24.477246

**Authors:** Weijie Yang, Jingsi Ji, Shuyang Ling, Gang Fang

## Abstract

Orthology prediction is challenging yet rewarding. Orthologs are the cornerstone of almost all comparative genomics studies. Dozens of ortholog resources have been available and broadly used over the past decades. However, the inconsistency between these resources has drawn growing concerns, especially when more proteomes are available and ortholog databases expand. It is no longer easy to decide which ortholog database to use, and it becomes necessary to compare conclusions based on different resources. We are presenting here a metric to assess ortholog functional consistency. Using this metric, we built a network connecting proteins based on their functional similarity. We then detected network communities as ortholog groups, and each protein in our ortholog group inherited the network degree centrality. By benchmarking Quest for Orthologs (QfO) and some representative ortholog resources, we concluded the degree centrality could serve as the index for the reliability of functional consistency. The numerical nature of degree centrality could also open a door for quantitative study in pan-genome and other comparative genomics studies.

## Introduction

To appreciate biological data, people must transfer knowledge from one species to another since most of the knowledge directly acquired from wet-lab experiments stems from a small group of model organisms. On this ground, according to the “ortholog conjecture”^1,2^, orthology is arguably the most fundamental concept in biology. In the past decades, the problem of identifying orthologs has drawn tremendous efforts from biologists and data scientists. Dozens of computational methods and resources to infer gene orthology relationships are available^3^. These resources, in one respect, significantly facilitated biological research; however, new problems also emerged, as the inconsistency between ortholog resources is too large to ignore^4–6^. This inconsistency has cast doubts on the precisions of many studies, such as those conclusions drawn from various pan-genome projects.

It was recently reported that using ARS (Adjusted Rand Score) and other indices to compare ortholog groups (OG) of last eukaryotic common ancestors, the average concordance of popular ortholog resources between eggNOG^7,8^, OrthoFinder^9^, Broccoli^10^, and others, is barely about 0.52^6^. Such high inconsistency is a call for reflection: is this mainly a technical issue or biology?

About half a century ago, when Walter Fitch first introduced the concept of orthology, he defined orthologs as genes descended from common ancestors through speciation events^11^. In this paper, Fitch also perceived that speciation and duplication interplayed, changing gene evolutionary fate and biological function. Now we understand the enormous variances between genomes, and it is natural to hypothesize that gene duplication occurred at a very high frequency, possibly more often than speciation. Some evidence suggests that all vertebrates evolved through two rounds of whole-genome duplications, and about 20-30% of human genes are ohnologs^12^. In addition, Koonin *et al*., among others, discussed that horizontal transfer and gene loss further complicated the evolutionary events, which caused a more diversified evolution of biological functions^13^. It appears that the discussion of orthology always has two connotations in one context: biological function and phylogeny. Another question thus appears: If duplications and non-vertical inheritances dominate protein evolution, when we build function-oriented ortholog resources, should we always root our work in phylogeny?

As we previously reviewed^5^: extant well-known ortholog resources could be classified into two categories: one category roots in phylogenies, such as TreeFam^14^ and Broccoli^10^; and the others are clustering of proteins based on their similarities, such as COG^15^ and eggNOG^7^. Among the phylogeny-based ortholog resources, a recent work of Broccoli implemented a strategy by first building a phylogenetic tree using representative proteins and then assigning possible in-paralogs into corresponding ortholog groups^10^. Compared to other methods mainly focusing on the congruence between a gene tree and species tree, Broccoli is faster and possibly constructs a more inclusive resource^10^. Nonetheless, when using phylogeny-based resources to annotate a new genome, we need to assume there is no functional divergence within the tree of an OG.

There is a more direct logic toward biological function for the other category of ortholog resource, pioneered by BBH (Bidirectional Best Hit) and COG^16^. BBH relies on the assumption of “one protein, one function” for most genes. E.g., a pair of mutually most similar proteins most likely perform the same particular biological function in two different species^5,16^, and this assumption has two essential premises: “function by single gene” and “function required in both species”^5^. But, of course, evolutionary events such as sub-functionalization and neo-functionalization could break the BBH premise. In such cases, a pair of mutually most similar genes may not have the same function. Besides, nascent gene duplication would harm the completeness of BBH ortholog groups^5^. Koonin and other pioneers noticed these issues from the beginning. The first version of COG was clustering of BBH triangles instead of single BBH^16^, and the following resources like eggNOG and OrthoMCL^17^ each had their adaptations to deal with the issues mentioned above^8,17^.

Nevertheless, these clustering-based methods all worked on partitioning one protein similarity matrix to identify communities. This partitioning strategy also drew some concerns discussed in our previous review^5^. The main question came from the selection of species, which could change the matrix’s community structure. Another concern was whether selective pressures on different OGs in the same matrix are distinctive enough to detect communities.

We investigated the above questions from a biology function point of view and designed a derived protein similarity network rooted in one assumption: protein sequence evolution is not continuous. This assumption is inspired by some recent works studying possible evolutionary dimensions of protein sequences^18,19^. In short, we assumed within the same ortholog group, amino acid substitutions could occur and accumulate, which doesn’t change the function of this ortholog group. So, within an OG, we could assume that protein sequence evolution is continuous; In contrast, when functional divergence occurs and new OG emerges, proteins in different OG should have enough amino acid mutations to allow a functional evolution. Thus, we proposed a non-continuous hypothesis regarding protein sequence evolution across other OGs.

To explore this non-continuous hypothesis, we leveraged a critical yet common understanding learned from systems biology^20,21^, such that there cannot be too many genes exerting the same function in one genome for most biological functions. Accordingly, we designed a simple 2-dimensional plot. Briefly, we chose some top similar proteins from a selection of fully sequenced species for each protein. For example, we took the top 10 most similar proteins from each of 300 selected bacterial genomes for a bacterial protein A. We assumed, in most species, there should be just one or two exerting the same biological function as protein A, and most other chosen proteins should belong to several divergent functional groups. We plotted these 3000 proteins on a 2D plot measuring their relative sequence similarity and length difference to protein A (Fig. 1A). For convenience, we defined protein A as a “seed”. Not surprisingly, in the 2D plot, we observed a large cloud of proteins overlaid and congested in a long and narrow area far away from the seed protein (Fig. 1A). This group of proteins was defined as “noises”, which are the mixture of several ortholog groups. In addition, a prominent group sat between the seed and noises, which we defined here as “signals”. This signal set is the group of proteins hypothesized as functional consistent orthologs to the seed. We reported the 2D plots for all over 2 million seed proteins in this paper, showing a pervasively clear distinction of signal and noise.

**Figure 1.**
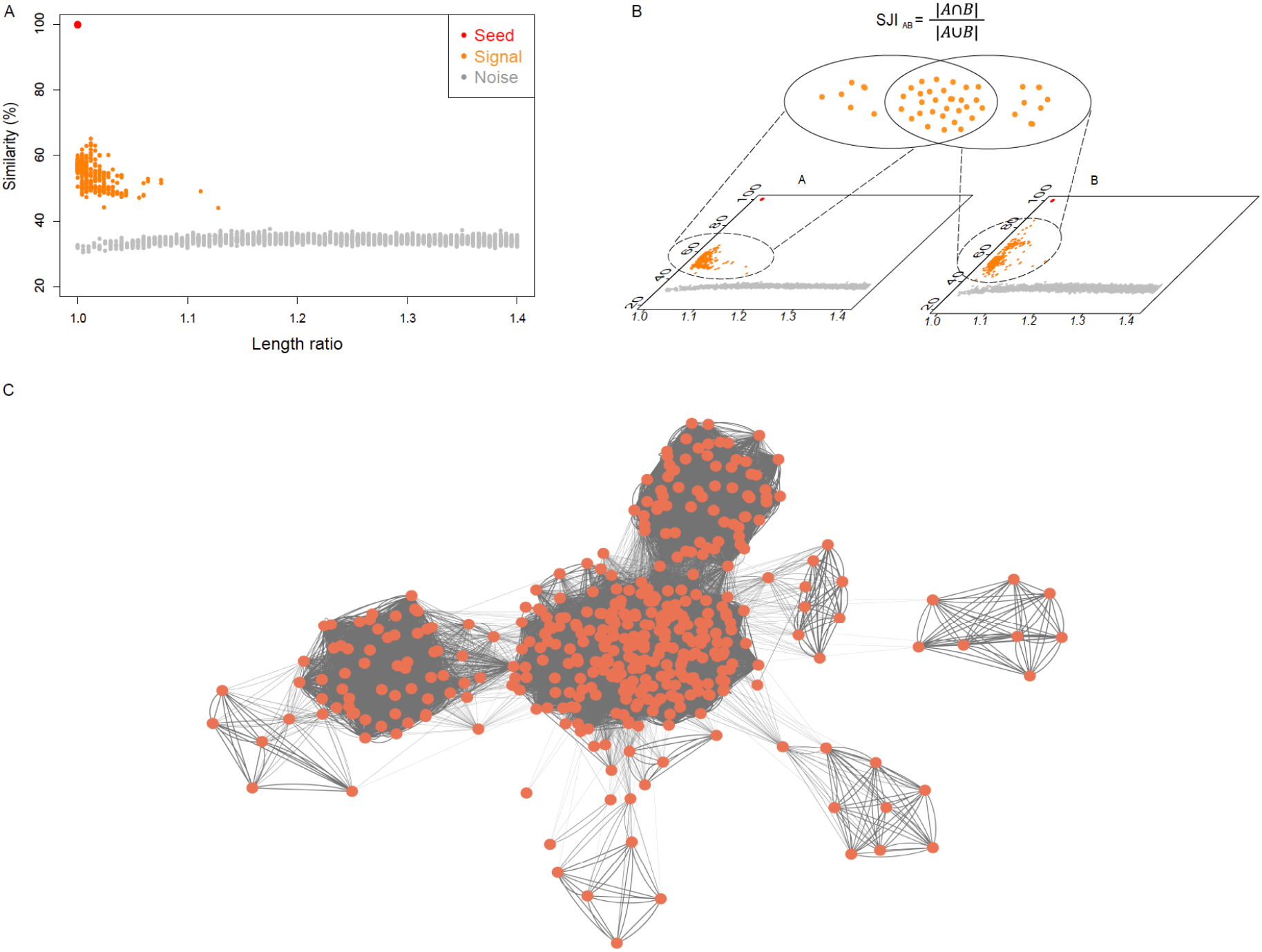
Overview of the workflow A) 2D plot displays relative coordinates of top similar proteins from different proteomes to a seed protein. The x-axis is the length ratio of proteins to the seed, using the longer length divided by the shorter. All similar proteins considered on a 2D plot have an upper limit of 1.5 in length ratio. The y-axis is protein similarity between proteins to the seed, calculated using opscan^38^. A spectral clustering algorithm separated the signals from the noises, colored in orange and grey. B) A schematic diagram shows Signal Jaccard Index. SJI describes the overlap of signals of two seeds, measured using the size of the intersection divided by the size of the union of two signal sets. C) A partial view of the derived network. Nodes are seeds, and edges are weighted using SJI.

If two seeds have the same biological function, we further assume the overlap of their signals should be high. Hence, we used the overlap of two signal sets, measured by the Jaccard Index, as the metric to assess the functional similarity between two seeds. Given the metric, we could build a derived network connecting all seed proteins, in which edges between proteins were weighted using this Jaccard Index. We constructed two such networks, one for bacteria and one for eukaryotes. Considering the high duplication rate^22^ in the eukaryotic genomes, instead of the top 10 from bacteria, we selected the top 50 most similar proteins for each seed so that enough functionally diversified noise proteins are on the plot to make the signals distinct.

With this network, we exploited the Louvain algorithm^23^ to detect communities as ortholog groups, benchmarked our OG to QfO^3^, and received a very high PPV (Positive Predictive Value) of 0.96. Additionally, network properties were reserved in each OG and on each protein. The most exciting finding is that using network Degree Centrality (DC) assigned to each protein, we could explain the inconsistency between primary ortholog resources such as TreeFam, eggNOG, Broccoli, OrthoFinder^9^, and SonicParanoid^24^. For proteins with high DC, these five independent resources tend to group proteins in the same OGs, showing a better consistency; those with lower DC show a opposite trend. DC analysis also revealed how the size of OG impacts the agreement of ortholog assignments. We suggest that DC is a compendious feature describing the reliability of protein functional consistency, and DC makes it possible to do the pan-genome studies quantitatively. We also discussed the potential evolutionary significance of the vast amount of low DC proteins and a few implications from the network topology.

## Results

### The clear-cut boundary between signal and noise in the 2D protein similarity plot

We obtained 1,231,088 bacterial and 889,930 eukaryotic proteins from the selected proteomes (Methods). Each of these proteins was a seed, and for which we looked for a vast number of candidate proteins from different genomes and plotted their relative similarity to the seed on a 2D plot (Methods). The vast number of candidate proteins guaranteed that there should be both functional consistent orthologs for each seed, if the seed was not a species-specific singleton, and remote homologs with diversified biological functions on the exact figure. Then a question left is whether the functional consistent orthologs, termed here as signals, are mixed with or separated from the noises, which refer to the overnumbered functional diversified remote homologs?

To investigate the above question, we examined the 2D plots, and found that distinct demarcations between signals and noises were consistent in all over 2 million plots. Figure 2AB shows two seed proteins as examples. One is the human olfactory receptor 52K1, reportedly subject to “unusually rapid evolution”^25^. Another is *ugpE* from *E. coli*, a transmembrane component in the ABC transporter complex, one of the largest protein family which is suggested to play an accelerating role in evolutions across all life domains^26^. We used these two proteins as examples for their fast evolution and the large protein family size: the number of their signals cannot be small, and their noises should be closer to signals. Fast evolving and large protein family size yielded a higher chance for the noises mingling with the signals. However, density plots (Figure 2AB) show apparent valleys between the signals and noises. And such observable valleys are ubiquitous, and all 2D plots are available at the database www.protdc.org we developed for this paper.

**Figure 2.**
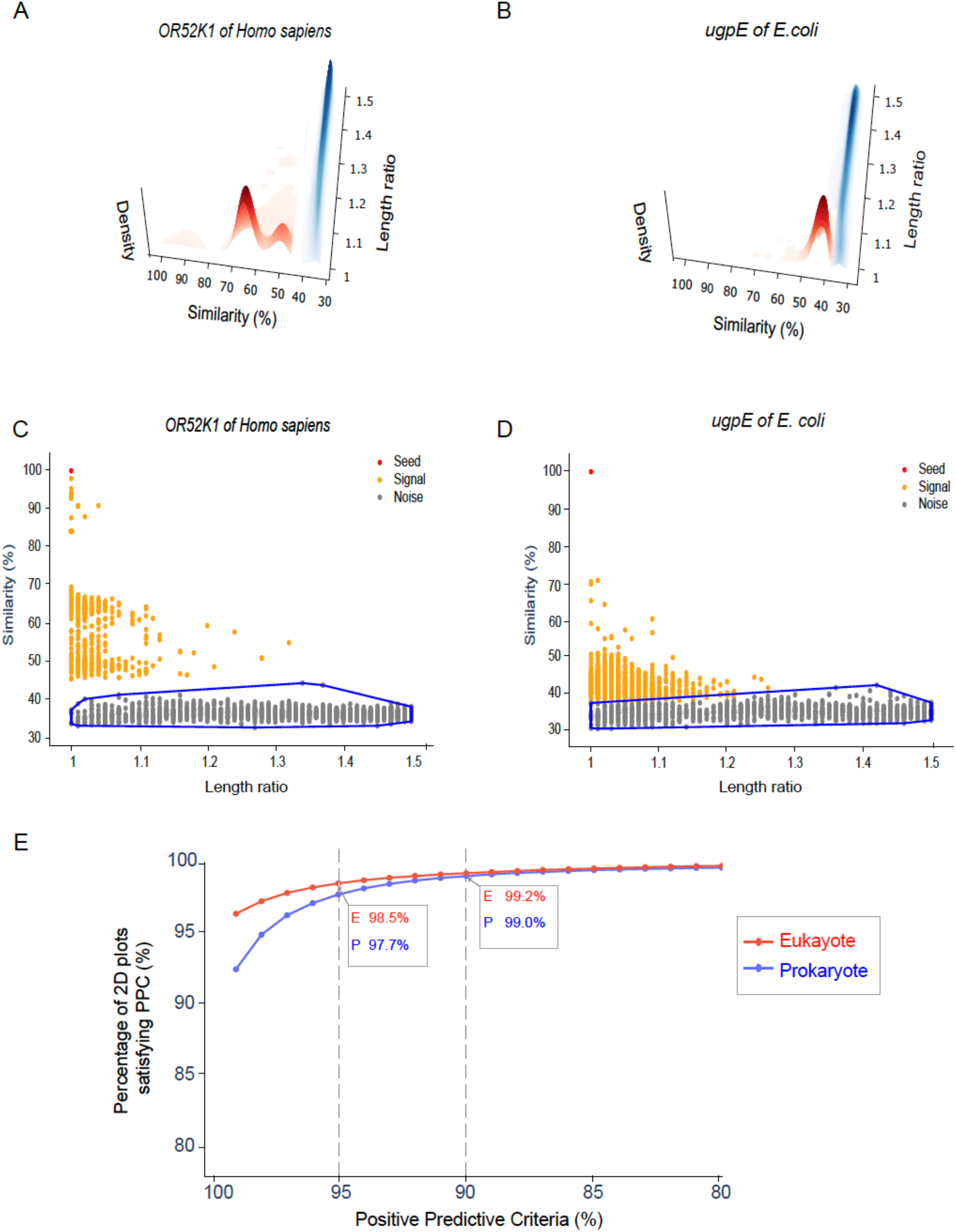
Non-continuous protein sequence evolution across OG A) A density plot shows the density of signals and noises of the human olfactory receptor OR52K1. Density is based on kernel estimation of kde2d R package^45^. B) Density plot of another example. The seed protein, *ugpE*, is a transmembrane component of the *E. coli* ABC transporter. C) A convex hull encircles noises of *OR52K1* drawn by the qhull algorithm from the SciPy package. D) Noise convex hull of *ugpE*. E) Evaluation of signal precision rate. This figure summarizes signal PPV for the whole over 2M seeds. Signal PPV on each 2D plot was defined as the percentage of signals floating above the noise convex hull. The x-axis is the critical values, and the y-axis is the percentage of 2D plots satisfying each PPV criterion.

We then deployed a spectral clustering algorithm^27^ to separate signals from the noises. Next, we drew a convex hull to encircle the area of noises. Higher the percentage of signals floating above this convex hull (Figure 2CD), better precision the signals; and this allowed us to estimate the signal’s PPV (Positive Predictive Value, see Methods). Fig 2E presents the overall PPV for eukaryotic and prokaryotic seeds under different criteria. It shows that in over 90% of all 2M plots, PPV is 100%, e.g., no signal falling into the noise convex hull; and 98.5% and 97.7% of 2D plots from eukaryote and prokaryote, respectively, satisfy PPV ≥ 95%.

We further evaluated the BLAST E-value of the seed-signal pairs. We took each seed as the query and BLAST against each proteome. Then we retrieved the E-values between all seed-signal pairs. 208M prokaryotic and 47M eukaryotic seed-signals pairs had E-values less than 10^−6^, accounting for over 97% of all seed-signal pairs. Interestingly, 150M prokaryotic and 46M eukaryotic seed-noise pairs also had strong E-values of less than 10^−6^, despite the apparent gaps between signals and noises (Supplementary Figure 1). We suggest this as a warning against using E-value or some arbitrary similarity cutoff to select ortholog candidates. The gaps between noises and signals reflect biological information between whole-genome contexts; but arbitrary BLAST E-values, though significant, make researchers blind to such gaps.

Based on the ubiquitous gaps between signals and noises and the high signal PPV (Figure 2E), we argue that a protein follows a non-continuous sequence evolution when OG is diversified and biological function evolves. With the Pfam database^28^ and many other studies working on multiple sequence alignment, it is well noted that certain amino acid sites are more conserved than others. Some recent work further revealed mutual constraints between such conserved sites, which were more related to protein function and determined protein fitness and stability^18,19,29^. The constrained amino acid sites cannot be the same in proteins with different functions. Because it is natural to understand that if one significantly mutates, the other constrained amino acid sites “optimize” in response, so the protein function evolves. Apart from the constrained sites, many other sites are subject to more independent and less limited mutations. These less biased mutations cause continuous sequence variations within an OG, which explains the variance within signals. And the chances for those less constrained sites across OGs being the same should be improbable. This explains the ubiquitous existence of a “valley”, but not a smooth and monotonic decreasing surface from the peak of “noises” to the area of “signal points” on the 2D plot (Figure 2AB).

### The metric and derived protein similarity network

With signals isolated from the noises, we could investigate the overlap of signals of different seeds. Suppose there is independent evidence supporting functional consistency between the seeds with the same signal sets; in that case, we have further justifications of assuming signals from the 2D plot as ortholog candidates.

Signal Jaccard Index (SJI) was used to measure the similarity between signals of different seeds (Methods). Supplementary Figure 2AB summarizes the SJI for prokaryotic and eukaryotic seeds. We identified 208M pairs with the median of SJI as 0.70 for prokaryotic proteins and 48M pairs of eukaryotes seeds with the median of 0.63.

We understand SJI as an extension of BBH. As we introduced, BBH directly connects pairs of proteins from two genomes with the same biological function. It considers the entire gene context from two complete genomes, but BBH finds a pair of genes based on gene-to-gene mutual identification; it cannot deal with recently duplicated genes with the same function or sub-/neo-functionalized genes belonging to diversified OGs. SJI could break these limitations. As long as enough noises are included, the gap between signals and noises appears. Recently duplicated genes with the same function should be expected in the signals, and the sub-/neo-functionalized genes, if they are not transient in evolution, and diversified OGs are formed and inherited by enough species, should be piled up in the peak of noises.

After obtaining SJI between all pairwise proteins, we built up a network connecting all proteins simultaneously. Network is one excellent way to integrate and purify biological information. Reliable biological information could be detected from network topologies^30^. Proteins with consistent high SJI due to strong evolutionary pressures will be grouped in communities. Special proteins with multifunctionality due to domain variations, would probably be the hub nodes with high betweenness. Supplementary Figure 2C-H summarizes the initial network consisting of proteins as nodes and SJI as weighted edges.

### Ortholog groups detection

Low SJI between some proteins was inevitable, making almost all the proteins connected in the initial network. However, the community structure in the network was evident, which allowed us to apply a gradual zoom-in strategy to identify ortholog groups (Methods). We used weighted community density≥0.25 as the clustering convergence threshold (Methods). This generated 73,503 eukaryotic and 72,541 prokaryotic OGs. This threshold can be adjusted. We tested a range from 0.1 to 0.25. The lower threshold, the larger but less precise OGs. After benchmarking QfO and other resources, we concluded 0.25 is the optimized threshold based on the following concordance comparisons between different ortholog resources.

Since our OG was detected as a community from the network, the degree centrality (DC) was assigned to each protein. DC was the sum of SJI from one protein to its neighbors and normalized by OG size (n-1). DC reflects how well a protein’s signals overlapping with neighboring proteins. To evaluate the stability of network structure and the reliability of protein DC, we made a bootstrap analysis. We randomly removed 10% of the proteins, followed the same procedures and thresholds to determine OG and protein DC; we repeated this process 1000 times. We compared the bootstrapped OGs to the original results using ARI (Methods), and this reported an average ARI of 0.93; the average correlation coefficient between bootstrapped DC to the original one was 0.85. This bootstrap analysis suggested a very stable network structure and protein degree centrality. Details of the bootstrap analysis are in Supplementary Figure 3.

All OGs can be downloaded from www.protdc.org.

**Figure 3.**
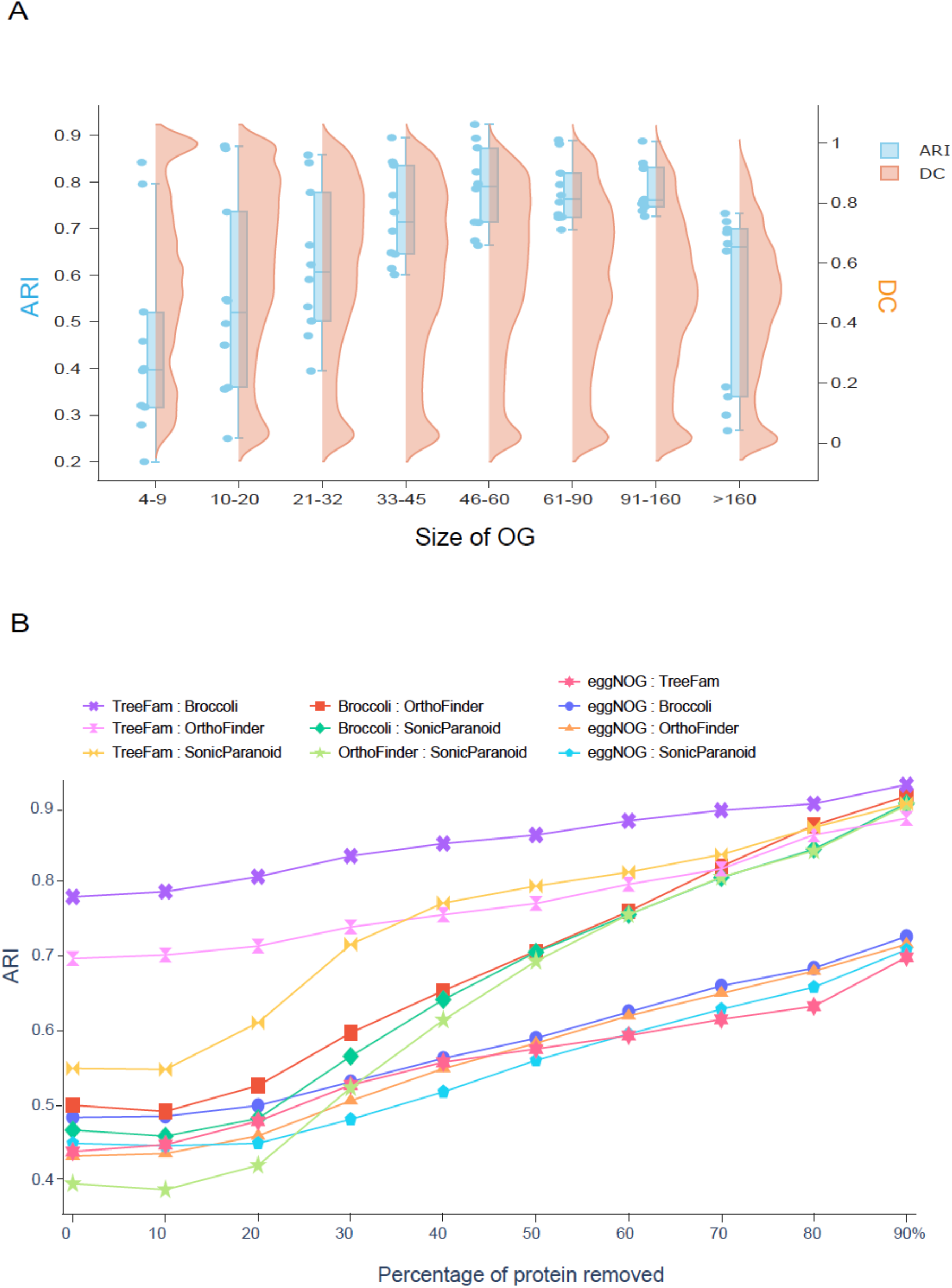
Protein degree centrality and the inconsistency between ortholog resources. A) OG size vs. Distribution of DC vs. ARI. We binned OGs according to their size. The x-axis is the size of OGs. The first bin includes over 103K proteins, from small OGs containing 4 to 9 proteins, and the rightmost is 86K proteins, from OGs containing at least 160 proteins. In between, these bins have from 80K to 100K proteins. The orange curve is protein DC distribution. DC is normalized 0 to 1 using OG size. In the left two bins (size <= 20), there are apparent biases toward larger DC, attributed to the smaller OG size. The blue boxplot describes the concordance between TreeFam, eggNOG, eggNOG, Broccoli, and OrthoFinder using ARI on the corresponding protein groups. Numerical details of the blue boxplot are in Supplementary Table 1. B) Concordance of ortholog resources improves in response to the increase of DC. 494K proteins from OG size >20 are the data subject to analysis in this figure. We organized this data into ten groups. Each group gradually removed 10% of the lowest DC proteins from left to right. For example, the leftmost group contains all 494K proteins, the second has 445K, and the rightmost group comprises about 49K proteins with the highest DC values. Concordance between ortholog resources on each group was calculated using ARI. Concordance between every pairwise ortholog resource shows a monotonic increasing trend.

### Benchmark ortholog group to other orthology resources

QfO is a service to evaluate ortholog prediction quality developed by a consortium of experienced scientists^3^. A frequent application of this service is to assess the concordance between the developer’s ortholog pairs and curated QfO references of SwissTree and TreeFam-A^3^. We leveraged this service and built our OG based on the QfO 2021 version of 48 eukaryotic proteomes. Unfortunately, QfO doesn’t have enough prokaryotic proteomes. We obtained 367 bacterial proteomes evenly distributed on the Greengenes^31^ phylogenetic tree from EMBL-EBI (Methods). Therefore, we could only benchmark TreeFam using our eukaryotic OGs. Supplementary Figure 4 summarizes detailed benchmarking results. We received a high precision such that PPV was 0.956, but the recall was mediocre, and TPR was 0.518. Most other ortholog resources on Supplementary Figure 4 showed TPR from 0.6 to 0.7. However, as discussed above, protein degree centrality can be used as an index delineating ortholog reliability. After removing proteins with DC<0.1 (accounting for 9.4% proteins submitted to QfO), PPV (0.958) almost didn’t change; TPR dropped 6.2% to 0.486. That said, removing low degree proteins caused an increase of “false negative” in TPR. In addition, considering the large numbers of seed-noise pairs with strong BLAST E-values discussed above, we suspected that a significant number of seed-noise pairs were related to our mediocre recall.

Accordingly, we studied the impact of gradually removing low degree proteins on the concordance between several ortholog resources, including TreeFam, Broccoli, OrthoFinder, eggNOG, and SonicParanoid. These representative ortholog resources utilize different algorithms. TreeFam is the curated reference in QfO and is reputed for its high quality. Broccoli and OrthoFinder represent phylogeny-based OGs. eggNOG represents clustering-based OGs and is also known for its high citations. Finally, SonicParanoid is included in this experiment for its genealogical relationship with InParanoid^32^, which identifies orthologs by more direct reasoning on biology function. Before the detailed examination, we first looked at the relationships between OG size, distribution of degree centrality, and background concordances among the five representative ortholog resources. The concordances were calculated using both ARI and AMI (Methods), which are highly consistent, and the numerical results are in Supplementary Table 1. Figure 3A shows an increasing trend of ARI among ortholog resources when OG size grows from 4 to 160. Since we obtained OGs following a gradual zoom-in process, the smaller OG size indicates proteins’ periphery location in the derived network. Figure 3A thus presents a coincidence between network periphery location and weaker ARI. We argue this is because the biology information from a low-density periphery network structure might be challenging to capture, and the biology information from a large OG is rather more accessible by different algorithms since the large OG was endorsed by a cohort of proteins sharing significant SJI. However, in Figure 3A, for proteins belonging to the largest OGs (size > 160), the concordances dropped, which means various resources had difficulty reaching an agreement on assigning orthologs belonging to fast-evolving and large protein families. Furthermore, Figure 3A shows a bias towards high DC in smaller OGs (size <=20). Thus, we only considered the intermediate- to large-sized OGs in the following experiment.

This experiment is to study the inconsistency among ortholog resources. We examined changes of orthology concordances in response to changes of protein degree centrality. We first pooled proteins from different OGs (size > 20) together, generating a dataset over 494K eukaryotic proteins. We sorted these proteins according to their DC and extracted ten groups shown on the x-axis of Figure 3B, and each group had 10% of low DC proteins removed from left to right. The curves in Figure 3B show the trends of ARI between ortholog resources on the ten groups. We can see a monotonic increase of concordances between all pairs of resources when gradually removing low DC proteins. For example, on the rightmost of Figure 3B, when removing 90% of low DC proteins, the ARI between TreeFam and three other resources increased to 0.85~0.90. The recent paper^6^ found high ARI values like this only on the 125 manually curated OGs. But in Figure 3B and Supplementary Table 2, the high ARI values were reached when considering over 1600 OGs covering 20K~45K proteins (Supplementary Table 2). AMI had higher raw values but the same trend as ARI (Supplementary Table 2). Supplementary Figure 5 provides additional evidence on DC’s consistency with overlaps from different ortholog resources, e.g., proteins from the intersection of more ortholog resources, higher the DC values.

A concluding lesson from this experiment is the power of degree centrality. The above results show we could obtain “high-quality” data sets comparable to manually curated OGs using DC to “purify” ortholog groups from various algorithms. The high-quality data, of course, is not from wet-lab experiments but from DC analysis which is supported by concordance from independent ortholog resources. Concordance is often the evidence commonly used in gene annotation^4^, and we argue high orthology concordance on high DC proteins is demonstrated in this experiment.

## Discussion

This study centered on using the SJI metric to measure protein function similarity. As discussed above, we regard SJI as a metric inspired by BBH. We followed the same logic by leveraging the information carried in the whole gene repertoire. But instead of finding directly 1:1 mutually identified gene pairs, we split the process into two steps. We first distinguished signals from noises, which generated a 1:n relationship between a protein and its signals; next, we used SJI to measure the overlap of “n” between two proteins, which gave a numerical weight in their similarity. This weight also carries sequence variation information between two proteins. For example, if two proteins have different domains, they should convey a low SJI. The derived network further integrated biological information carried by SJI from single proteins into a community in an extensive system. The network community structures reflect the evolutionary results on proteins from different genome contexts. Therefore, back to the example of different domains, we believe those proteins should have a certain distance on the network, belonging to diverse communities.

Next, we suggest DC as a compendious feature describing the reliability of protein function. Biology function is essentially implemented via the recruitment of protein sequences. It should be pointed out that orthology deals with comparing sequences and information carried by gene repertoires, not directly biology functions. It is on the basis of “ortholog conjecture”^1^ that orthology bridges biology functions obtained from limited wet-lab experiments to the enormous genome data. Biology function can be promiscuous^33^, moonlighting^34^, and even fuzzy^35^, but this isn’t equal to poor concordance in orthology, although orthology provides only an indirect view of biological function. Thus, understanding the inconsistency of various orthology prediction algorithms and finding a way to work it around becomes an important question. And our answer is DC. DC is a network feature rooted in SJI, reflecting biology information carried in the whole gene repositories. DC is independent of a priori threshold; as long as genomes used to obtain DC are not too similar in their genome context, DC should be reliable. And we believe DC will be particularly useful in pan-genome studies, where the inconsistency of different ortholog resources is well noted and poses severe problems in the field^36^. And the numerical nature of DC will allow quantitative analysis of genes instead of classifying them into two groups, the pan- and core genomes.

Besides, we also noticed many proteins with low DC or clustered in very small OG. A rough estimation (OG size < 20 or DC < 0.2) is that about 40-50% of proteins are low DC proteins. We assume these proteins should be subject to future work. In the future, when more genomes are included, those large OGs identified in this work might be stable, but the amount of low DC proteins might be ever-increasing. If the low DC proteins are due to transient evolutionary events or on the path to disappearing, they would have low eigenvector centrality and other genres of centralities. However, some promiscuous ancestral proteins^33^ with multiple domains could also link to two or several clusters in the network, and we guess the betweenness and eigenvector centrality might be higher in such a case. We know the theory of “one protein, one function” doesn’t apply to all, but how many proteins have just one major function, and how many do not? Where do the non-duplicated moonlighting proteins locate in the network, and how many functions are connected to the moonlighting proteins^34^? Based on this derived network, many protein evolutionary questions might be investigated quantitatively by in-depth topological analysis and studies on protein’s various genres of centralities.

## Online Methods

### Eukaryotic proteomes and selection of bacterial genomes

Quest for orthologs (QfO) is a community-developed web service to facilitate benchmarking ortholog prediction methods. To leverage this service, we downloaded the 2021 version of 48 eukaryotic reference proteomes from the QfO website^37^ to build this function-oriented protein evolution network. Since QfO doesn’t include many bacterial genomes, we downloaded bacterial genomes from EMBL’s European Bioinformatics Institute (EMBL-EBI) website. Fully sequenced bacterial species are highly biased with pathogens and model organisms. To reduce sampling bias, we downloaded the 2013 phylogenetic tree based on 16s rRNA from Greengenes^31^ and cut the tree into 400 clades to ensure the evolutionary distances among these clades were roughly even. We then selected one genome out of each unbiased phylogenetic clade, resulting in a total of 367 bacterial proteomes (some clades had no fully sequenced genomes). All bacterial and eukaryotic proteomes used in this study are at www.protdc.org.

### 2D plot and spectral clustering to detect signals

Each protein is a seed. The top 10 and top 50 most similar proteins to the seeds from bacterial and eukaryotic genomes, respectively, were selected using the tool of opscan^38^. Opscan measures protein sequence similarity using the FASTA algorithm and BLOSUM62 matrix^38^. We restrained the top similar proteins with protein length difference less than 1.5 (using the length of longer one divided by the shorter), mitigating the workload due to multiple domains variance. We then plotted each seed and its similar proteins on a 2D plot (Figure 1A), with the x-axis being the protein length ratio ranging from 1 to 1.5 and the y-axis being protein sequence similarity.

On each 2D plot, we deployed a spectral clustering algorithm^27^ to separate the signals from noises. This algorithm’s major steps are as follows:

Input: two-dimensional points of size *n*, where *n* is the number of proteins.

I. Zero mean normalization on both dimensions.
II. Construct a similarity graph G by measuring the Euclidean distance between each point, and get the adjacency matrix A and degree matrix D.
III. Compute the random walk normalized Laplacian matrix *Lrw*=*D^−1^L*=*I-D^−1^A*
IV. Determine the number of clusters *k* that maximizes the eigengap.
V. Find the smallest *k* eigenvalues and the corresponding eigenvectors *x1, …, xk* of *Lrw*.
VI. Denote U as a matrix containing the eigenvectors *x_1_, …, x_k_* as columns, its size is *n* × *k*, and *n* is the number of points.
VII. Use the K-Means algorithm on U to partition the data points into *k* clusters *C_1_, …, C_k_.*

Output: Clusters *C_1_, …, C_k_*.

Due to our setup, the largest cluster with the highest data points density was always composed of noises. Sometimes, there were a few smaller clusters to the right of the largest one. Because of the exceeding protein length difference, these clusters were noises by default. All other clusters floating above these noises were signals.

We further drew a convex hull encircling the noises to evaluate our clustering performance. We defined PPV (Positive Predictive Value) of signals as the percentage of signal points above the noise hull (Fig. 2CD). This convex hull was drawn using the qhull algorithm from the SciPy python package^39,40^. We also examined the BLAST E-value in between seed-signal vs. seed-noise.

### Derived protein similarity network

Due to biological information carried in complete genomes’ gene repertoires, signals are distinguished from noises. Each protein is a seed, and seeds with the same biological function should better overlap in their signals. Thus, we used the Jaccard similarity coefficient as an index to measure the functional similarity between seeds. Fig 1B illustrates the calculation of the Signal Jaccard index (SJI), which was the intersection of two signal sets divided by the size of their union. SJI also served as the weight of edges between all pairwise seeds, and thus a function-oriented network connecting all proteins was automatically constructed (Fig. 1C).

### Ortholog group detection

With the network, we used the Louvain algorithm^23^ to detect communities as ortholog groups. The essence of the Louvain algorithm is to maximize the modularity of communities^41^. Due to inevitable low-weighted edges, the initial network was huge, and applying Louvain once was not enough, as some large sub-networks with apparent clusters therein were common. Therefore, we adopted a step-by-step zoom-in approach to repeatedly use the Louvain algorithm until the community satisfies a density threshold. Density D considers edge weights,

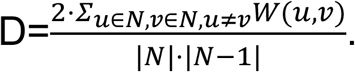

Where N is nodes in a community, W is the weight between node *u* and *v*. Below are the steps of ortholog group detection process, and the convergence threshold was density≥0.25.

Input: A weighted protein functional similarity network.

I. Calculate network density D. If it is larger than 0.25, stop the network cutting process and consider the current network as an OG; otherwise, continue to the next step.
II. Apply the Louvain algorithm to cut the current network.
III. For each sub-network, update edge weights by considering only nodes (seeds) included in the sub-network *per se*.
IV. Repeat from the first step for each sub-network.

### Benchmark orthology resources

To evaluate our OG, we did two categories of benchmarks. One benchmark was to refer to QfO. We followed the same procedure QfO processed eggNOG OG^3^. We exploited the ETE toolkit v3^42^ and followed the suggested eggnog41 workflow to construct a rooted phylogenetic tree for each OG^42^. Following this workflow, the tree balances efficiency and accuracy. Then using the species overlap algorithm^43^ from the ETE3 package, we obtained ortholog pairs. Since the bacterial proteomes used in our study were different from QfO, we processed only eukaryotic OG and submitted the results to benchmark QfO TreeFam-A ortholog pairs.

Benchmarking QfO centered on the phylogenetic consistency between our OG versus TreeFam-A. Still, the metric and network in this paper focus on leveraging biological constraints born in the whole-genome context. We assigned in-paralogs with the same biological function in the same OG. Accordingly, we compared our OG to TreeFam, eggNOG, Broccoli, OrthoFinder, and SonicParanoid using another two indices, Adjusted Rand Index (ARI) and Adjusted Mutual Information (AMI)^44^. Both of them are commonly used in comparing clustering results. ARI reflects the consistency of clusters based on pairwise counting, and AMI is rooted in Shannon information^44^.

## Source code availability

Scripts used in this work are available at https://github.com/FangLabNYU/protdc.

## Acknowledgments

We would like to thank Prof. Antoine Danchin, Dr. Jungseog Kang, and Dr. Jianfei Zhao for valuable discussions and comments on this article.

## Authors’ contributions

GF conceived and designed this study and wrote the manuscript. WY and JJ performed the analysis. SL advised on the algorithm and wrote the manuscript. All authors have read and approved the submitted manuscript.

---

Supplementary Table 1. ARI and AMI on protein sets sorted by OG size. This table contains the numerical ARI values plotted in Figure 3A.

Supplementary Table 2. ARI and AMI on protein sets sorted by protein DC. This table contains the numerical ARI values plotted in Figure 3B.

Supplementary Figure 1. Eight examples show significant seed-noise similarity measured by BLAST E-value (<10^−6^). BioProject ID or NCBI taxonomy ID is in the panel title. The green spots in the noise cloud have significant seed-noise E-values of < 10^−6^.

Supplementary Figure 2. Numerical overview of SJI and the initial network. Panel A and B show the boxplots of SJI for the prokaryotic and eukaryotic seeds; Panel C and D are degree distribution of the entire network; Panels E-H are correlations between four representative centralities, degree, closeness, betweenness, and eigenvector on nodes of four large sub-networks. We used sub-networks since the initial networks were too large that the meaningful relationships between these centralities should thus be from local communities between neighboring structures. The yellow histograms show distributions of the four centralities in the sub-networks, and grey scatter plots with blue density contours present their relationships.

Supplementary Figure 3. Bootstrapping results of ARI and DC. Each bootstrap had 10% of proteins removed, and we repeated the bootstrap 1000 times. Correlations between bootstrapped and original OGs are shown on the left, and the right panel is the distribution of the correlation of protein degree centrality.

Supplementary Figure 4. A screen copy of the QfO benchmarking result. Our dataset is marked as “protdc”.

Supplementary Figure 5. Intersections of different ortholog resources vs. protein degree centrality. The QfO v2021 eukaryotic proteins were extracted from the five representative resources: TreeFam, eggNOG, Broccoli, OrthoFinder, and SonicParanoid. The x-axis is the number of resources agreeing on ortholog assignments. For example, the rightmost bin has 59K proteins that were in the same OGs in all five resources; and the leftmost bin contains 61K proteins grouped in only one resource; in between, 172K, 212K, and 185K proteins have 2, 3, and 4 resources, respectively, supporting their OG assignment. The y-axis represents the distribution of degree centrality of proteins in the corresponding bin. DC positively relates to the concordance of different resources.

## Notes

https://www.protdc.org

### Competing Interest Statement

The authors have declared no competing interest.

