## Supplementary figures and images for "A metric and its derived protein network for evaluation of ortholog database inconsistency"

### Supplementary Figure 1

#### Prokaryote

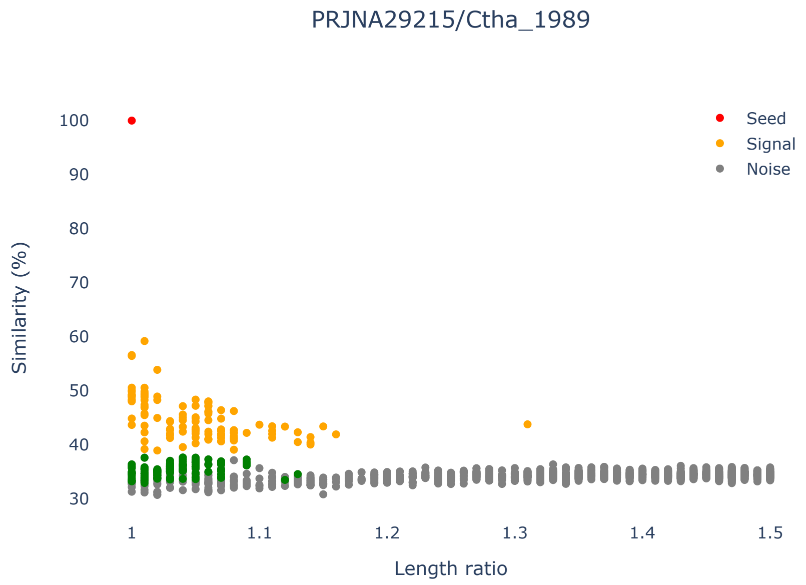


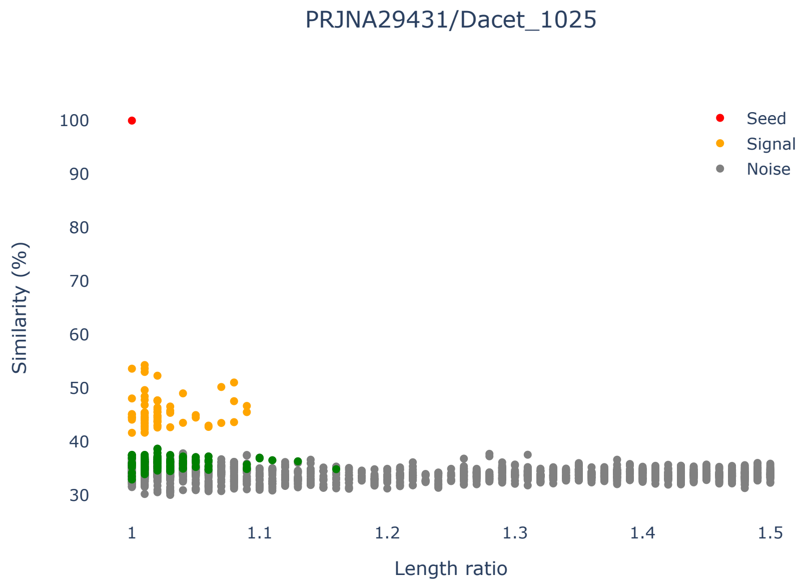


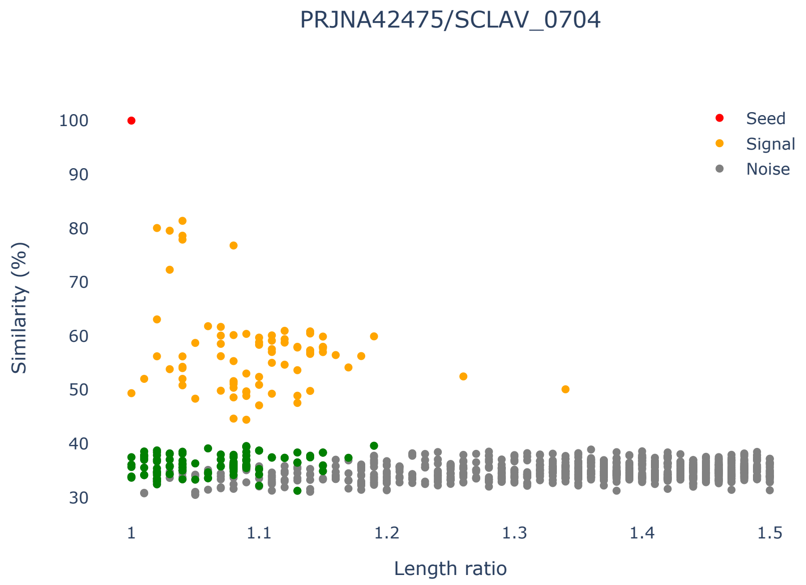


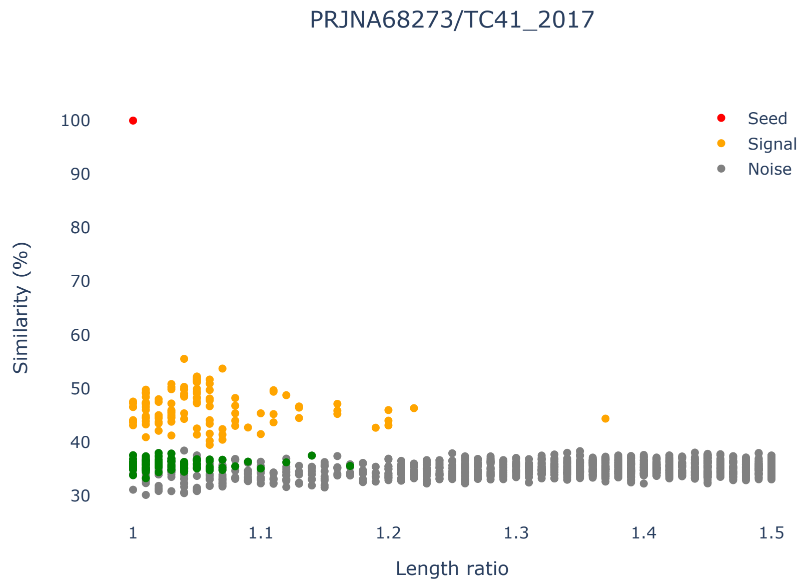


#### Eukaryote

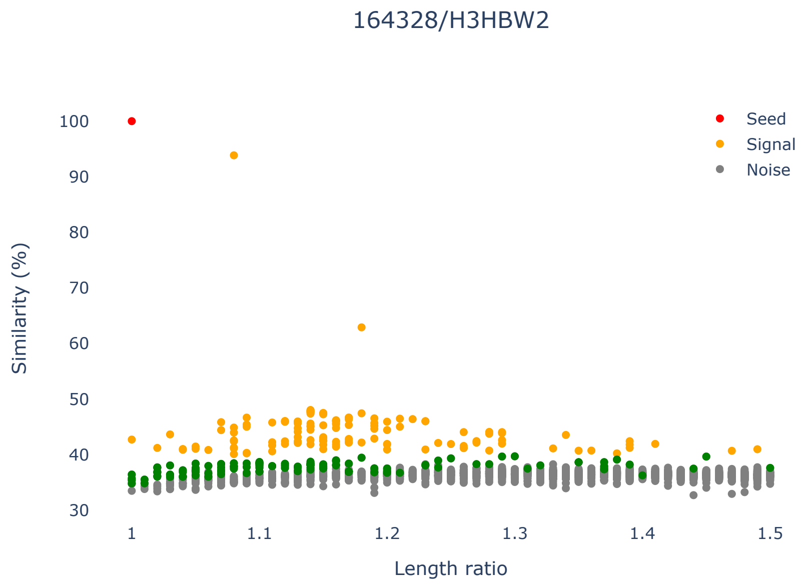


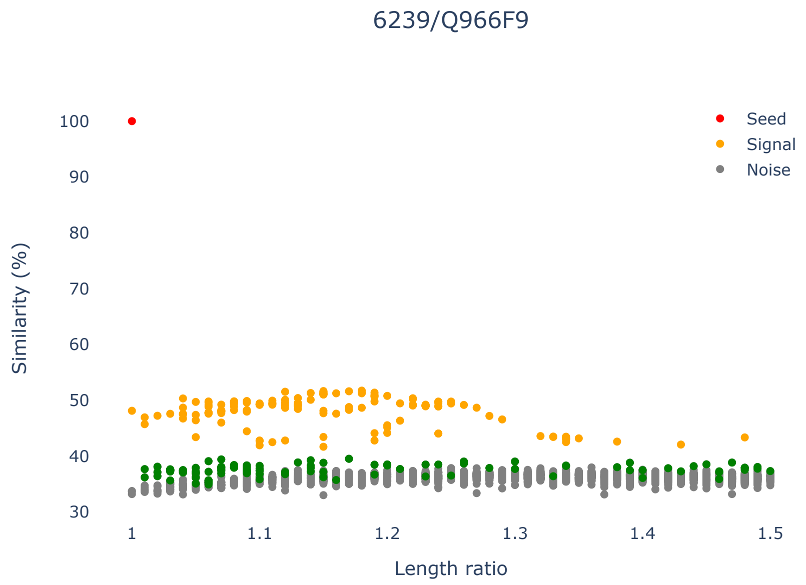


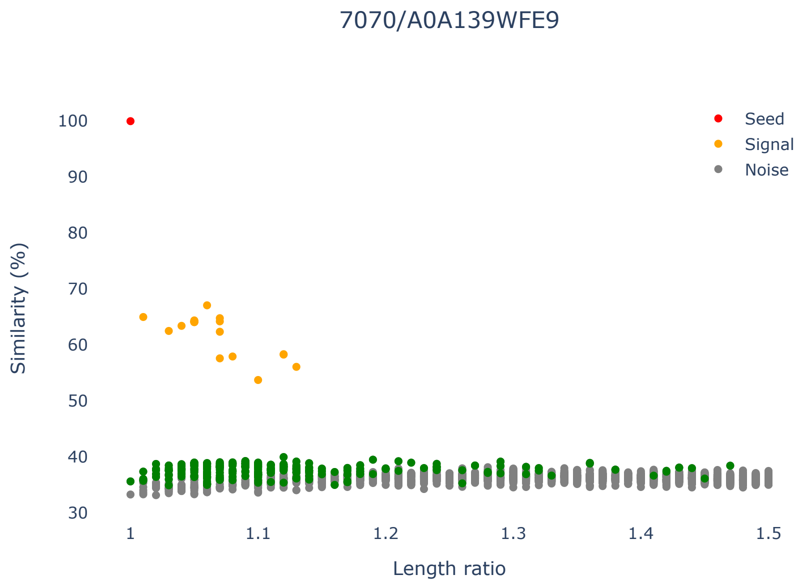


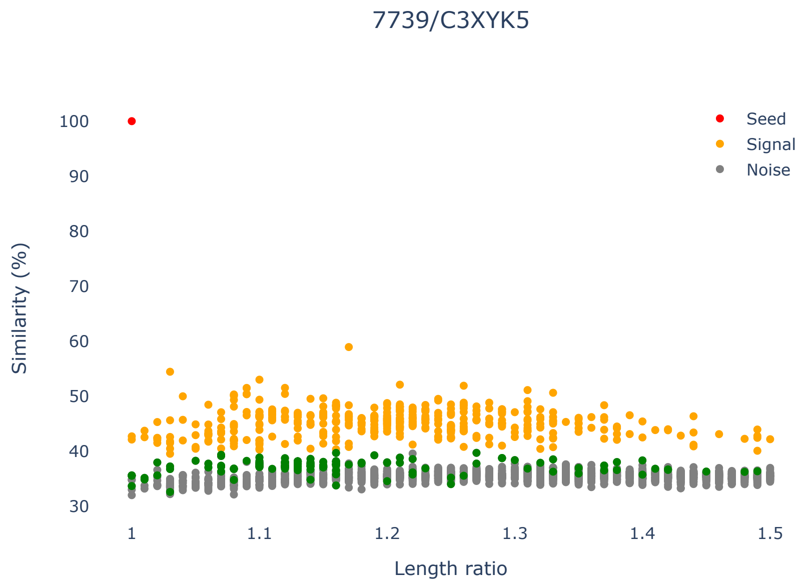

### Supplementary Figure 2

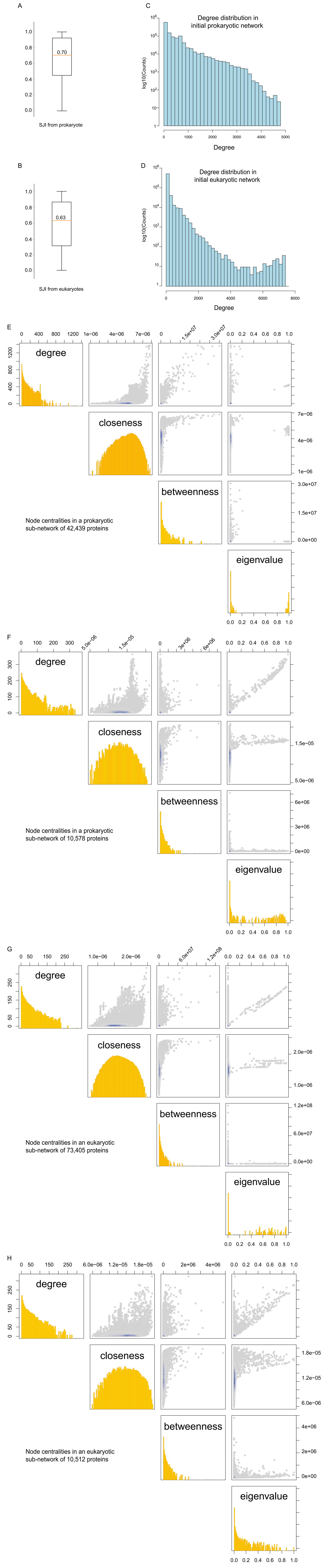

### Supplementary Figure 3

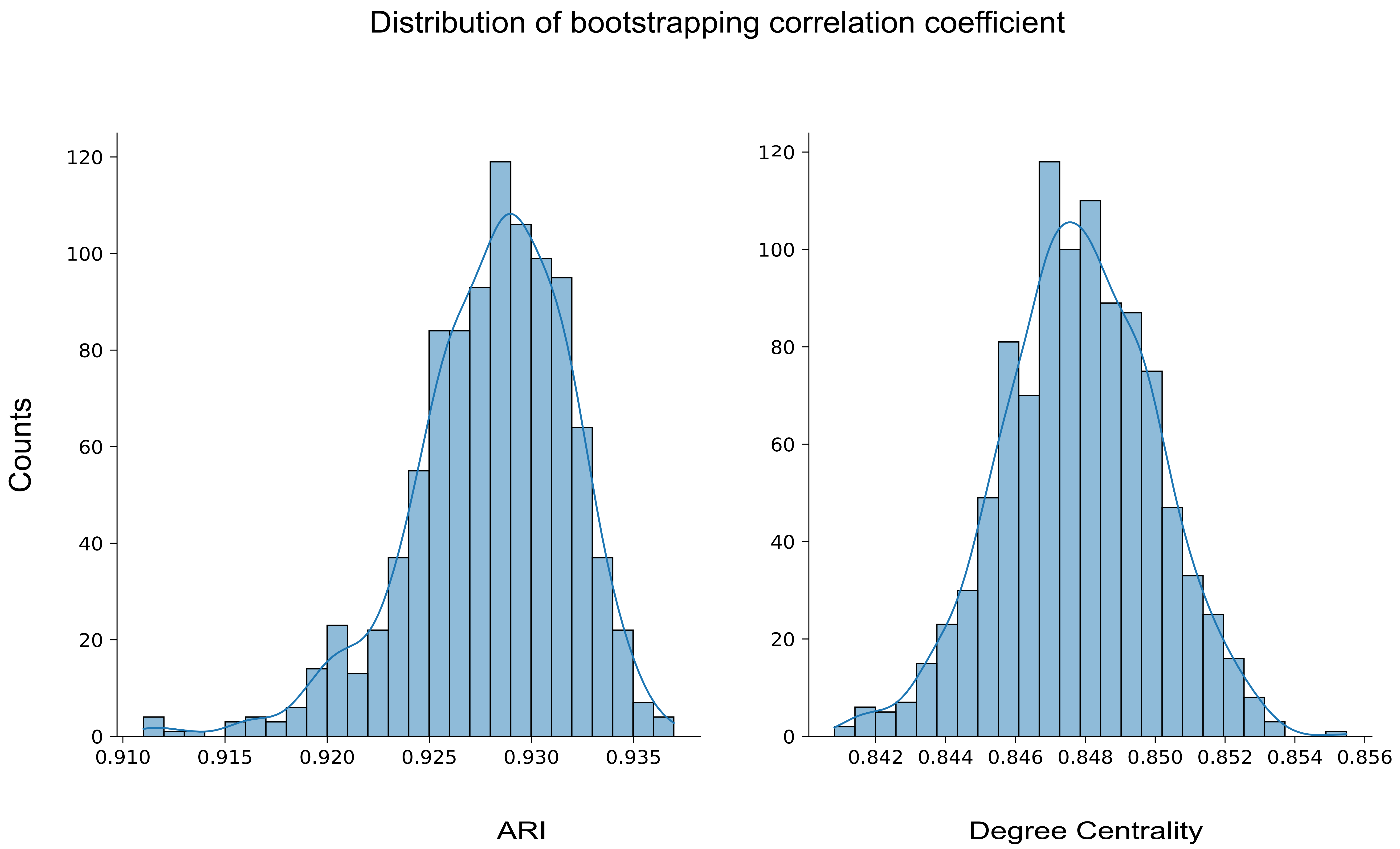

### Supplementary Figure 5

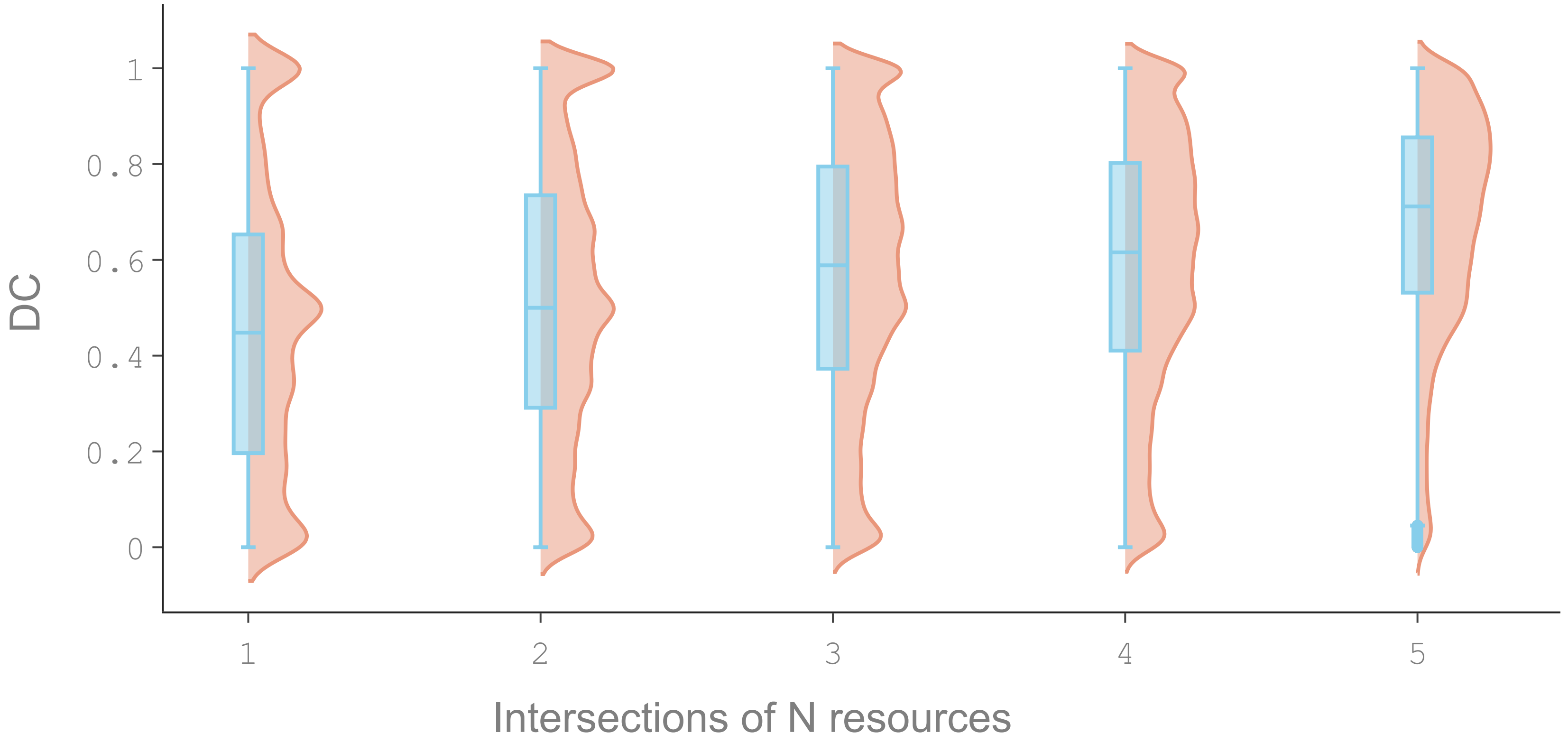
