## Supplementary Figure 4 for "A metric and its derived protein network for evaluation of ortholog database inconsistency"

### Challenge name: TreeFam-A - Agreement with Reference Gene Phylogenies: TreeFam-A

NO CLASSIFICATION

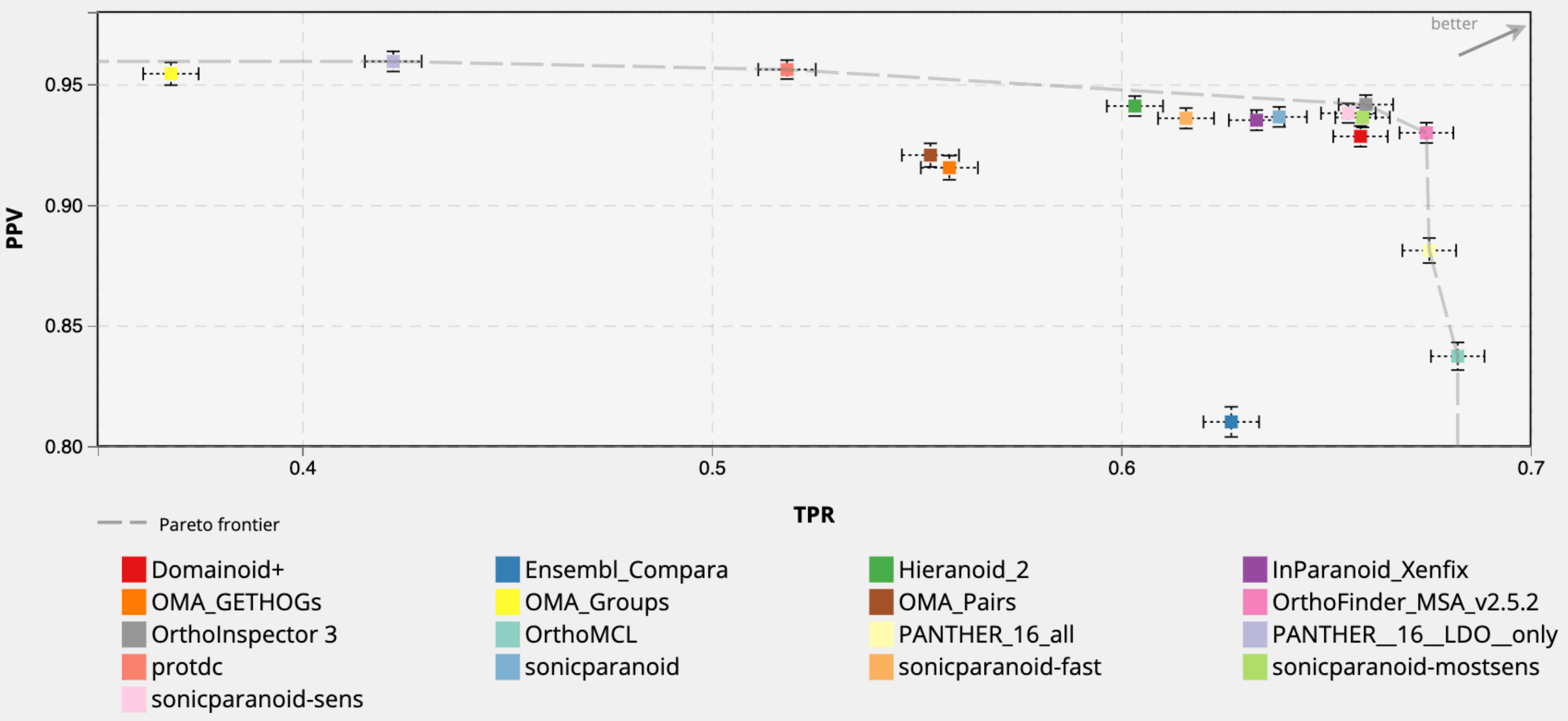
